# Evaluating the analytical validity of mutation calling pipeline for tumor whole exome sequencing

**DOI:** 10.1101/2022.11.17.516840

**Authors:** Chinyi Cheng, Jia-Hsin Huang, Jacob Shujui Hsu

## Abstract

Detecting somatic mutations from the patients’ tumor tissues has the clinical impacts in medical decision making. Library preparation methods, sequencing platforms, read alignment tools and variant calling algorithms are the major factors to influence the data analysis results. Understanding the performance of the tool combinations of the somatic variant calling pipelines has become an important issue in the use of the whole exome sequences (WES) analysis in clinical actions. In this study, we selected four state-of-the-art sequence aligners including BWA, Bowtie2, DRAGMAP, DRAGEN aligner (DragenA) and HISAT2. For the variant callers, we chose GATK Mutect2, Sentieon TNscope, DRAGEN caller (DragenC) and DeepVariant. The benchmarking tumor whole exome sequencing data released from the FDA-led Sequencing and Quality Control Phase 2 (SEQC2) consortium was applied as the true positive variants to evaluate the overall performance.

Multiple combinations of the aligners and variant callers were used to assess the variation detection capability. We measured the recall, precision and F1-score for each combination in both single nucleotide variants (SNVs) and short insertions and deletions (InDels) variant detections. We also evaluated their performances in different variant allele frequencies (VAFs) and the base pair length. The results showed that the top recall, precision and F1-score in the SNVs detection were generated by the combinations of BWA+DragenC(0.9629), Bowtie2+TNscope(0.9957) and DRAGMAP+DragenC(0.9646), respectively. In the InDels detection, BWA+DragenC(0.9546), Hisat2+TNscope(0.7519) and DragenA+DragenC(0.8081) outperformed the other combinations in the recall, precision and F1-Score, respectively. In addition, we found that the variant callers could bias the variant calling results. Finally, although some combinations yielded high accuracies of variant detection, but some variants still could not be detected by these outperformed combinations. The results of this study provided the vital information that no single combination could achieve superior results in detecting all the variants of the benchmarking dataset. In conclusion, applying both merged-based and ensemble-based variants detection approaches is encouraged to further detect variants comprehensively.

## 1. Introduction

High-throughput sequencing technologies have been integrating into clinical practices. The candidate genes panel-based sequence, whole-exome sequence (WES), and whole-genome sequence (WGS) are the most popular approaches. Recent studies[1, 2]have broadly demonstrated that the use of WES orWGS data can lead precise cancer treatment. Then, different treatments were given to the patients based on the cancer variant detected. Since the sequencing price has dramatically dropped over the past years, the ability to correctly identify the cancer variants is crucial for cancer genetic testing. A typical analysis pipeline includes the read alignment, variant calling, and variant annotation. The sequence aligner maps the sequence reads against the reference genome; the variant caller detects the variants from the aligned reads and outputs into the variant calling format (VCF) files. Nowadays there are many bioinformatic tools and tool combinations to achieve similar performance. However, previous studies have shown that the aligner and the variant caller could heavily influence the results of variant detection[3, 4]. Notably, understanding the performance of the tool combinations has been becoming an essential issue in the use of whole-genome sequences (WGS) or whole-exome sequences (WES) analysis on clinical actions. The conflicting results can dramatically affect the downstream clinical decision. Additionally, several studies indicated that different tool combinations often generate discordant variant detection results[5–10]. While some studies [5–10] have evaluated the effectiveness of the multiple combinations of aligners and variant callers on the variant calling analysis workflows, their focus has been primarily on germline variant detection by using different benchmarking datasets. It is lack of benchmarking dataset and tool comparisons. Therefore, this study aims to compare the performance of the different tools and their combinations.

Currently, there is no gold standard pipeline yet that benchmarks which methods work best in somatic variant detection, especially in tumor-only mode. In contrast to germline variant detection, the caller must be sensitive to somatic variants, which often have relatively low allele frequencies. On the other hand, the tumor-only mode is indeed desired by the clinic. However, the ground truth dataset for tumor-only mode mutation detection is extremely valuable. Until recently, the FDA-led Sequencing and Quality Control Phase 2 (SEQC2) consortium has released a benchmarking dataset for cancer variant detection in tumor-only mode[11]. Consequently, the evaluation of different somatic variant detection tools has become feasible.

In this study, we focused on the comparison of different combinations of aligners and callers. Many reputable aligners and callers were selected to evaluate the performances of cancer variant detection on the tumor-only benchmarking dataset. The aligners of BWA[12], Bowtie2[13], an Illumina DRAGMAP (https://github.com/Illumina/DRAGMAP), the build in aligner of the Illumina commercialize DRAGEN Bio-IT Platform (abbreviated as DRAGEN-A) (https://www.illumina.com/products/by-type/informatics-products/dragen-bio-it-platform.html), and a graph-based aligner HISAT2[14] were selected. The callers for cancer variant include GATK Mutect2[15], Sentieon TNscope [16], Illumina DRAGEN Bio-IT Platform build in caller (abbreviated as DRAGEN-C), and Google DeepVariant[17] were selected. Detailed information of all aligners and callers is given in Supplementary Table S1.

We reanalyzed SEQC2 benchmarking dataset and compared total of 18 combinations of aligners and callers for their performances on tumor-only mode in cancer single nucleotide variants (SNVs) and small insertions/deletions (InDels) detection. Our results provide critical information and evidence that no single combination produced superior results in detecting all the variants from the samples. In turn, the merged or ensemble-based approach, which combines results from multiple combinations, is most likely to improve the performance of variant detection in tumor samples.

## 2. Materials and methods

### 2.1 Benchmarking dataset

We downloaded the benchmarking dataset released by SEQC2 in 2021[11] (SRA accession: SRR13076390 to SRR13076398) and renamed as Sample1 to Sample9 in our study. The detailed information for this dataset is given in Supplementary Table S2. The truth VCF files and confidence regions (bed files) were downloaded from https://figshare.com/articles/dataset/Consensus_Target_Region/13511829. These FASTQ files were the results of the sequencing on a mixed cell lines set with different WES enrichment kits and libraries. This data set contains more than 40,000 variants including 41, 673 SNVs and 539 InDels variants, respectively. In these variants, there are 53.16% of variants whose variant allele frequency (VAF) is lower than 10%.

In this study, the SEQC2 released benchmarking dataset was composed of nine technical replicates by three exome kits including Roche, IDT and Agilent from three laboratories, respectively[11]. We downloaded FASTQ files by SRA tools with the RUN ID SRR13076390 to SRR13076398. These FASTQ files are the results of the sequencing on a mixed cell lines set with different WES enrichment kits and libraries. The truth VCF files and confidence regions (bed files) were download from https://figshare.com/articles/dataset/Consensus_Target_Region/13511829. This true positive variant set contains more than 40,000 variants including 41, 673 SNVs and 539 InDels variants, respectively. In these variants, there are 53.16% variants whose variants allele frequency (VAF) are lower than 10%.

### 2.2 Pipelines implementations

To evaluate the performances for each combination, we implemented the pipelines from FASTQ to VCF (see flowchart in Fig. 1). First of all, we applied five aligners, BWA, Bowtie2, DRAGEN-A, DRAGMAP and HISAT2, for mapping the input FASTQ files to the reference genome sequence, in here we used the hs37d5 as the reference genome. Then, forty-five (five aligners × nine samples) text-based Sequence Alignment Map (SAM) files were obtained. These SAM files were converted into the compressed binary version format files (BAM) by using the Sentieon built-in tools. The steps of the post-processing of BAM file, including sort, deduplicate, and local alignment, were done by using the Sentieon built-in tools. The detailed command lines were given in Table S2. Four variant callers, GATK Mutect2, Sentieon TNscop, DRAGEN-C and DeepVariant, were used to produce the result of variant detection for each BAM files. The compatible issue happened on the out BAM files by HISAT2 with the variant callers, DRAGEN-C and DeepVariant. As a result, in a total of 162 VCF files were obtained. The post-processing of these VCF files was conducted by a self-developed script to deal with the multiallelic problems in GATK Mutect2. GATK LeftAlignAndTrimVariants was used to normalize the variants and GATK SelectVariants was adopted to filter out the variants which were not in confidence regions. Finally, Haplotype Comparison Tools (https://github.com/Illumina/hap.py) and in-house developed programs were used to evaluate the performances the combinations.

**Fig. 1.**
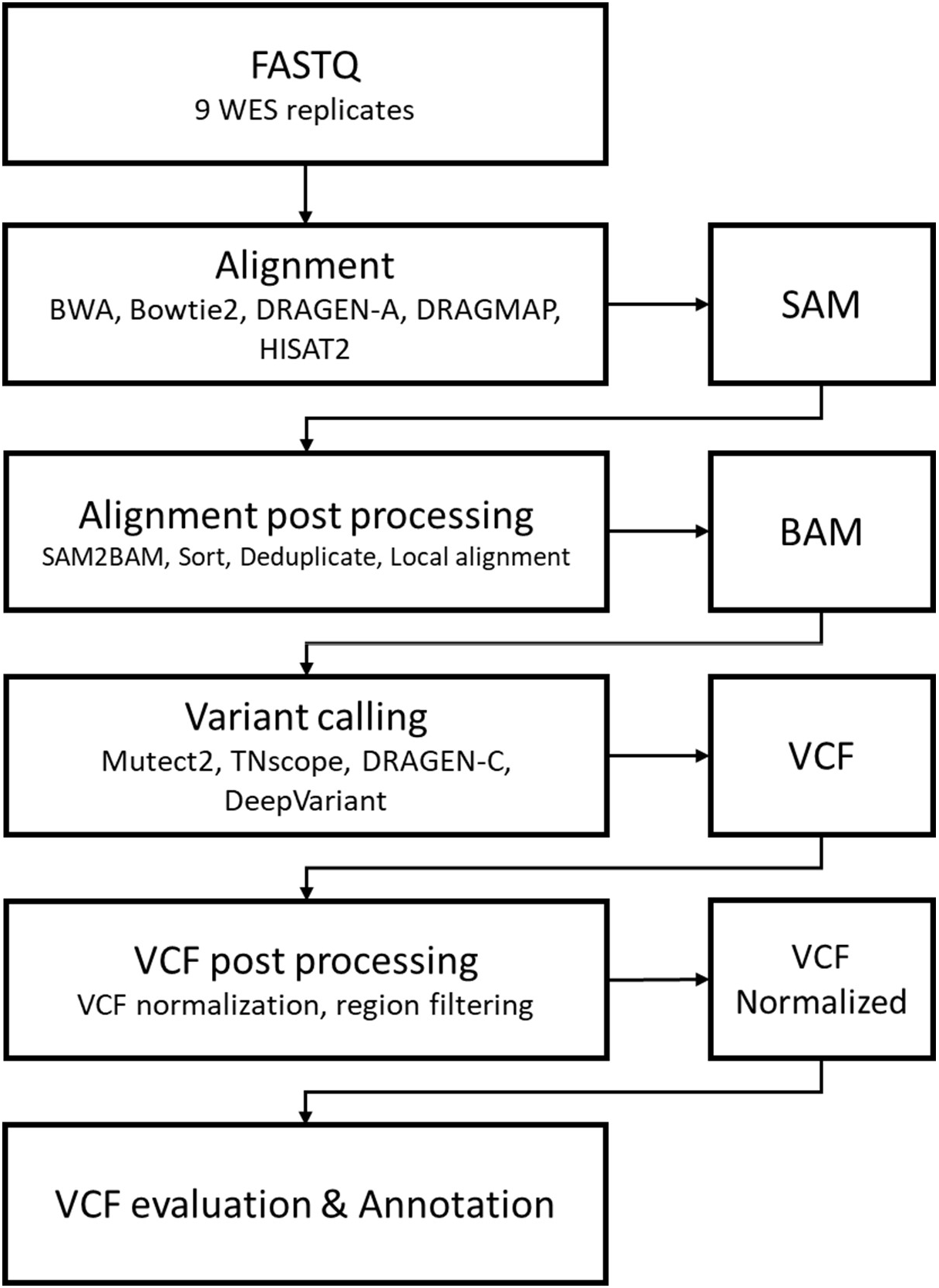
The flowchart of the benchmark analysis pipeline of different aligner and caller combinations.

### 2.3 Performance evaluation of variant calling pipelines

In order to evaluate the performance of every combination in different samples, we measured the

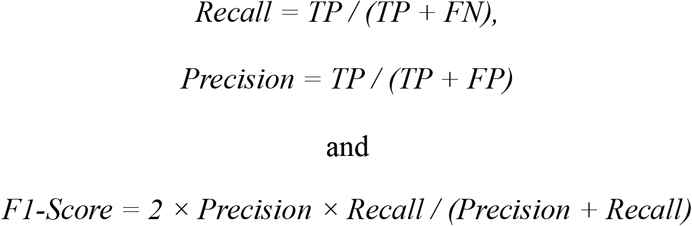

for every VCF output by utilizing Hap.py software from GA4GH [https://github.com/ga4gh/benchmarking-tools]. Every VCF output was used to measure the Recall, Precision and F1-Scores. An in-house pipeline which applied relational database to store the variants detected by each combination in different samples was developed. Finally, we utilized the in-house pipeline for some detail analysis and evaluations.

### 2.4 Computing environment and resources

All pipelines and sequencing data were run and stored on the high-performance computing (HPC) cluster computing environment within Taiwania 3 of National Center for High-performance Computing, Taiwan. In-house developed programs were used to measure the performances of each combination. For DeepVariant, we obtained the docker image of DeepVariant from https://github.com/google/deepvariant and installed on a Google Kubernetes Engine (GKE)-based HPC cluster computing environment with Taiwan AI Labs.

### 2.5 Cancer variants annotation and comparisons

## 3. Results

### 3.1 Results of detecting SNVs and InDels

For the comparison, we first focused on the F1-scores and total 18 combinations in all samples were shown in Fig. 2 (further details are given in Additional file 3: Table S3 and Additional file 4: Table S4). We separated the results into SNVs and InDels. As shown in Fig. 2A, the combination of DRAGMAP+DRAGEN-C showed the top performance with a mean F1-score of 0.98 in the SNVs detection. By contrast, the combination of bowtie2+deepvariant showed the lowest mean F1 score in the SNVs detection, The DRAGMAP+DRAGEN-C yielded best performance with a mean F1-score of 0.81 in the InDels detection (Fig. 2B). and in Sample2 was lowest than the other combinations and Samples in the InDels detection. The average of F1-Scores for every combination in different samples also shown that the combination of DRAGMAP+DRAGEN-C outperformed the other combination in both SNVs and InDels detections. For individual benchmarking dataset in the SNV detection, all the tool combinations yielded the best performance on Sample3 and the poorest performance on Sample4. And the mean F1-Score of Sample2 was the highest and Sample8 was the lowest in the InDels detection.

**Fig. 2.**
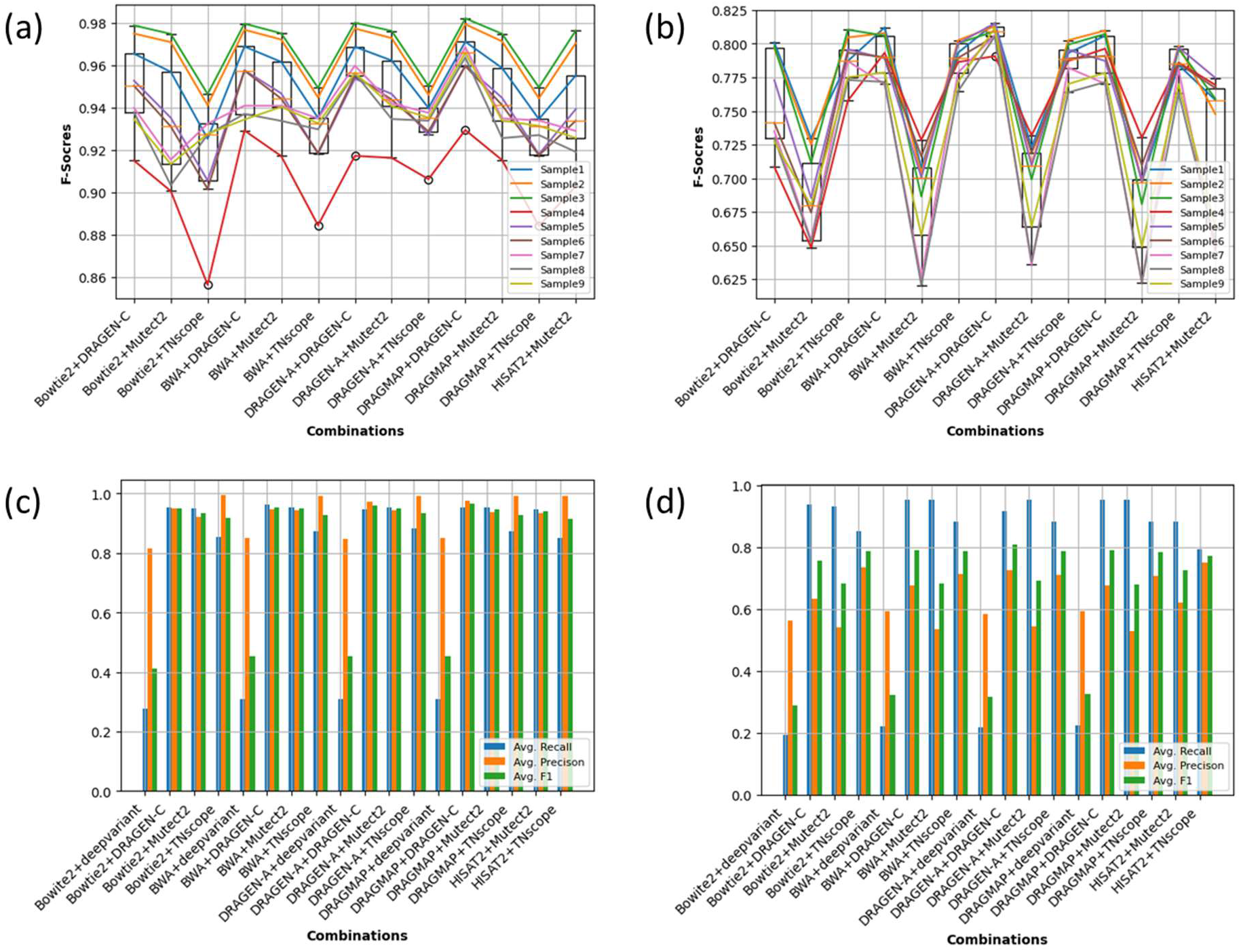

Next, we further analyzed the recall and precision of the combinations. The detailed results of the averages of the recall, precision and F1-Score for each combination in different samples in SNVs and InDels detections was shown in Fig. 2C, Table 1 and Table 2 (more detailed results are given in Table S3 and Table S4). Accordingly, we noticed that even though the combination of DRAGMAP+DRAGEN-C outperformed the other combinations in regard to the mean F1-score, the combinations of bwa+dragen and bowtie2+TNscope outperformed other combinations with better Recall and Precision in SNVs detection (Fig. 2D). In the InDel detection, the combinations of BWA+DRAGEN-C, HISAT2+TNscope, and DRAGEN-A+DRAGEN-C outperformed the other combinations, respectively, in the mean Recall, Precision and F1-Score. By contrast, the combination of bowtie2+deepvariant obtained the poorest performance in three metrics in the SNVs detection overall. In InDels detection, the combination of bowtie2+deepvariant obtained the lowest mean recall and F1-Score and dragmap+Mutect2 showed the lowest mean precision.

### 3.2 Performance in detecting SNPs and InDels at different

#### variant allele frequencies

One of the major differences between the germline mutation and somatic mutation detection is that the VAFs of the somatic mutation are always lower than the germline ones. Therefore, higher sensitivity is required for the somatic mutation detection pipelines.

In this section, the results of the averages of the numbers of the variants detected from all samples by each combination in different VAF in SNVs and InDels detection were shown in Fig. 3 (the detail results were given are given in Additional file 3 and Additional file 4: Table S3 and Table S4). The histograms of the VAF in SNVs and InDels of the truth VCF were also given as the references in the Fig.3 (a) and (b).

**Fig. 3.**
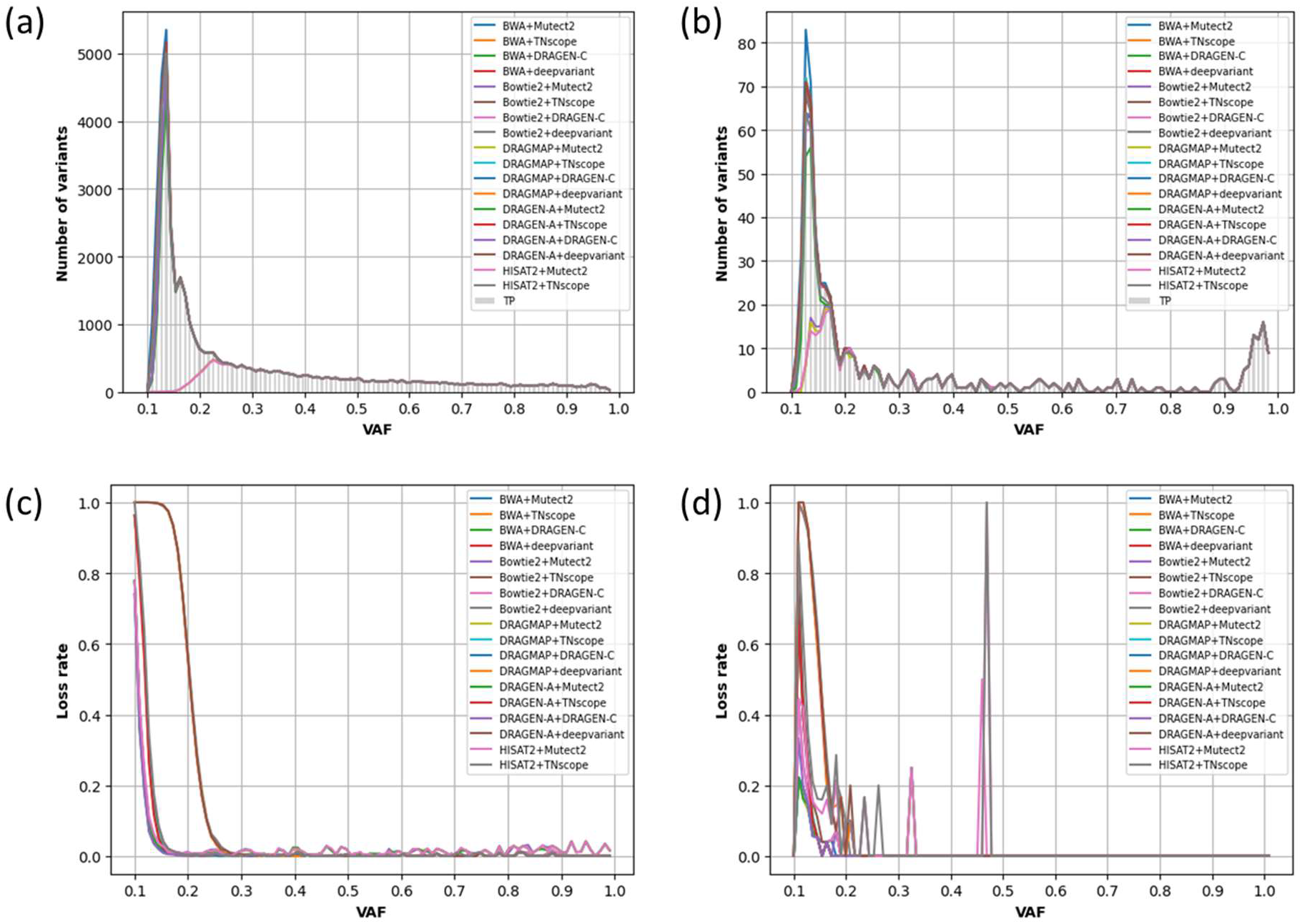

As the results are shown in Fig.3(a) and Table S4, most of the tool combinations demonstrated the abilities to detect almost all variants as VAFs were high (VAF >10%) in the detection of SNVs. However, those tool combinations showed poor detection on those variants with low VAFs (<=10%). Especially, the calling results produced by the combinations using DeepVairant as caller were unable to detect most variants with VAF less than 20%. In the results of the detection of InDels, as shown in Fig. 4(b) and Table S5, most of the combinations demonstrated the ability to detect most of all variants as VAFs were in high (VAF > 6%) and poor performance as the VAFs were in low (VAF < 6%). The results produced by the combinations whose caller was DeepVairant were approaching complete also lately, until the VAF over than 10%.

The results of loss rate for each combination in different VAF of SNVs and InDels shown in Fig. 5 (further detail results were given in Additional file Table S5 and Additional file Table S6). The loss rate was defined as

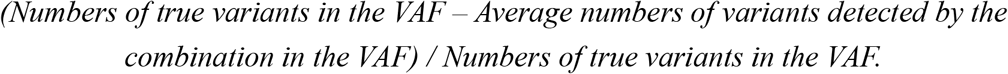

As the results show Table S5, the combination bwa+dragen produced the lowest average of the loss rate and dregen+deepVariant produced the lowest average of the loss rate in the detection of SNVs. On the other hand, in the detection of InDels, as shown in Table S6, the combination DRAGMAP+dragen produced the lowest average of the loss rate and bowtie2+deepvariant produced the lowest average of the loss rate.

To further understand the closing behaviors between the combinations, the raw data of the numbers of variants detected by the combinations in different samples and VAFs in SNVs and InDels detection were given in Table S7 and Table S8, respectively. We applied those raw data to the dimensionality reduction technique, t-SNE [18] and data visualization, and then generated plots, as shown in Fig. 4. These plots showed that the same caller produced the similar results on different samples.

**Fig. 4.**
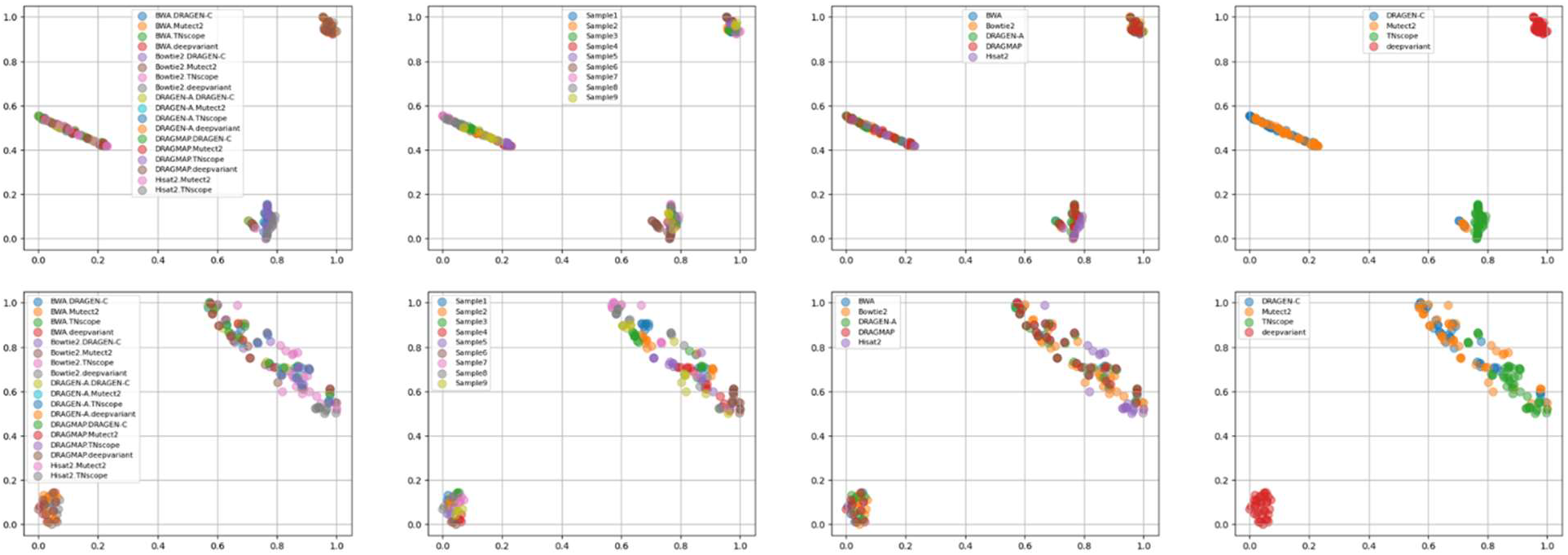

### 3.3 Performance in detecting InDels at different base pair (bp) length

**Fig. 5.**
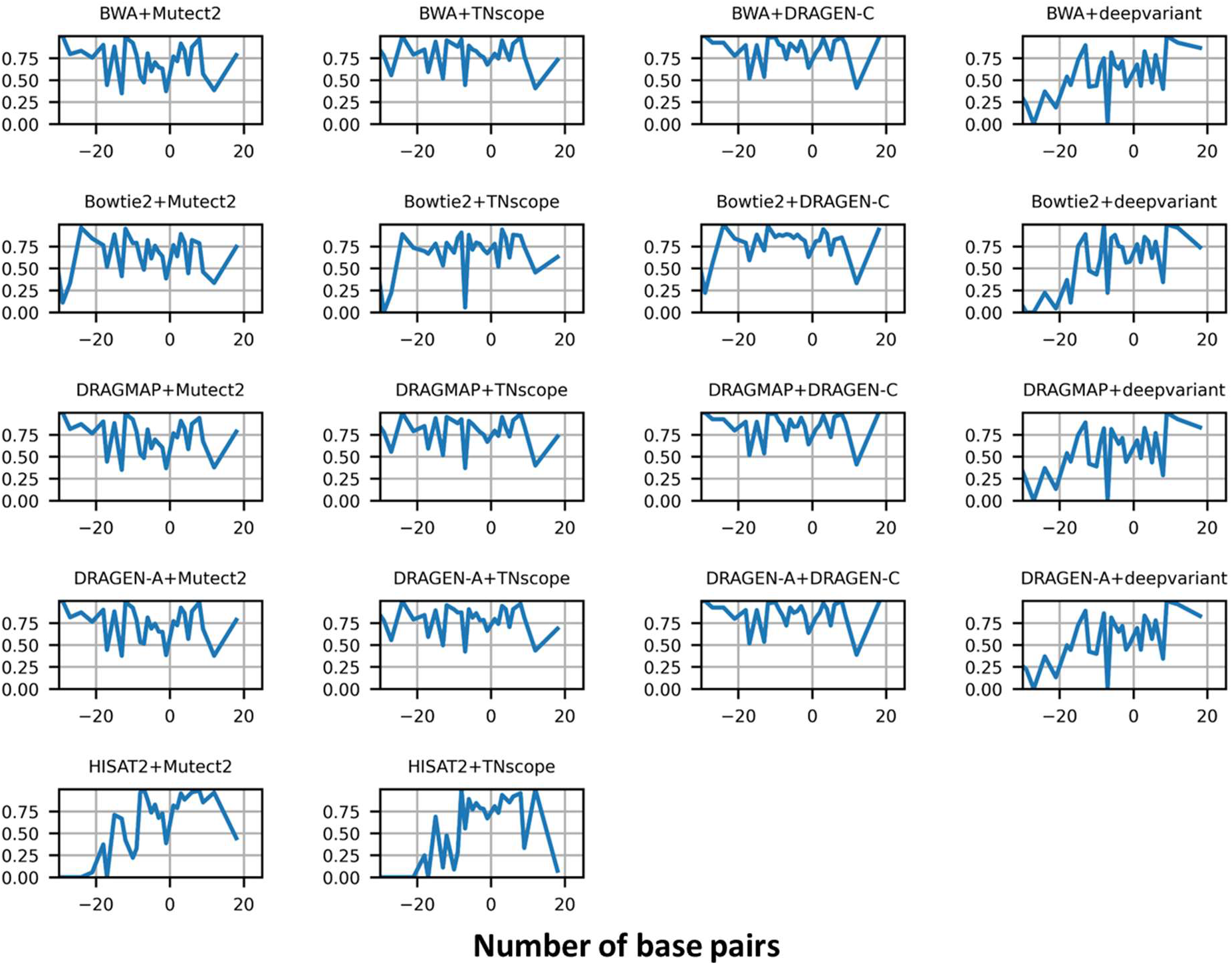

### 3.4 Shared variants in different combinations

Fig x and Fig x demonstrated the intersection of the variants by using different callers for each aligner in the SNVs and InDels detection, respectively. As shown in Fig. x, almost all aligners combined with the callers presented the almost same results, excepted from the results of the aligner HISAT2, in both SNVs and InDels detections. These results supported our finding in the section of ???, the aligners seemed did not influence the final results of variant calling obviously. We also noticed that even though aligners combined with the caller DragenC covered most of the numbers of detected variants. However, there were still a few variants which were only could be detected by the other callers. A critical issue is raised if these variants, which were not detected by DragenC, might be vital for identifying certain diseases. [we need an example] DeepVariant could not detect any new variant in the combinations whose algners were BWA, DragenA and DRAGMAP.

On the other hand, the venn diagrams presented the intersection of the variants by using different aligners for each caller in the SNVs and InDels detection, respectively (Fig. X). Although almost all callers combined with the aligners presented the almost same results, but the numbers of the common variants for all callers were lager than the ones for all aligners, obviously. We noticed that the results of the DeepVariant with the aligners showed poor detection on many variants that were detected with other caller tools. Specifically, there were 508 variants only could be detected by the combination with the Bowtie2 (Fig. X (d)).

## 4. Conclusions and future work

In this study, we evaluated the performances of 18 combinations of aligner and variant callers for somatic variants detection by using the benchmarking dataset released by SEQC2. A major difference with other previous benchmarking studies was that we focused on the usage of tumor-only model for variant detection. In fact, because for the clinical applications, the tumor and normal paired samples were always difficult to be obtained.

As the results shown in…different practices of wet lab techniques may influence the results of variant detection, even though the same pipeline was utilized.

Some callers performed the poor results of variant detection in low VAF like DeepVariant. The callers in the combinations may influence the results of variant detection in different VAFs.

The results of shared variants in different combinations shown that even if some combinations have demonstrated superiority in the F1-scores, but there were still a number of variants that which did not detect. Those number of the variants might be some important variants for some diseases.

That’s one of the reason some studies applied the merged or ensemble-based approach which majorly combines the results of variant calling from multiple variant caller or combinations to improve the accuracy of variant detection.

The results of this study provide supportive evidence that the merging the results of multiple variant caller or combinations could improve the results of variant detection by using single or few variant caller or combinations.

